# Hierarchical Encoding of Regulatory Mechanisms and Expression Syntax by a foundational genomic sequence-to-function model

**DOI:** 10.64898/2026.01.10.698352

**Authors:** Jiaqi Li

## Abstract

Deciphering the comprehensive regulatory rules encoded in the genomic sequence remains a central challenge in functional genomics, requiring a paradigm shift from descriptive annotations to a mechanistic understanding of the entire genome. Here, we introduce HERMES (Hierarchical Encoding of Regulatory Mechanisms and Expression Syntax), a framework that progressively defines a fundamental sequence vocabulary for the complex regulatory genome and parses gene expression syntax into transparent biological insights. Specifically, we trained a foundational sequence-to-function model on a massive compendium of 137,127 functional genomics profiles spanning diverse biochemical marks and cellular conditions, including DNA methylation, transcription factor binding, polymerase binding, histone marks, chromatin accessibility and RNA expression across various tissues, cell lines and cell types. By harnessing the high-fidelity sequence representations of HERMES, we established a context-specific regulatory vocabulary comprising 40 distinct sequence classes and fine-grained subclusters spanning the entire genome. We interrogated the sequence determinants of diverse promoters and tissue-specific enhancers, and further identified a housekeeping genes-specific promoter sub-class, driven by ETS, YY1 and CCAAT motifs. To further decode transcriptional regulation, HERMES was leveraged to predict cell-type-specific gene expression and the enhancer perturbation effects. Extensive evaluations confirmed that the cis-regulatory elements and their interactions were captured to predict gene expression underlying different cellular conditions, and simultaneously revealed divergent enhancer dependencies between housekeeping genes (HKGs) and highly variable genes (HVGs). Ultimately, we distilled the complex sequence-to-function model into a biologically interpretable rulebook of Enhancer-Promoter interaction grammar. Synthesizing all the insights, we propose a unified compatibility model, where HKGs utilize a strong-promoter architecture for high-output expression, while HVGs depend on a context-dependent syntax driven by compatible promoters and enhancers. In summary, HERMES bridges the gap between predictive modeling and biological mechanism, transforming sequence representations into a comprehensive functional encyclopedia and a quantitative grammar of the complex regulatory genome.

## Introduction

Deciphering the language of genome sequence remains a fundamental challenge for functional genomics^1^. Historically, genes have been regarded as the major functional units of the genome, which are spatiotemporally expressed in distinct cells to establish diverse biological phenotypes^2,3^. With the completion and analysis of the human genome, researchers revealed that genes comprise only a minor fraction of the genome, and most of the genome regions consist of non-coding sequences, which orchestrate the spatiotemporal regulation of cell-type specific gene expression^4,5^. To understand the function of the non-coding regions, large-scale consortia like ENCODE and Roadmap Epigenomics have comprehensively delineated the putative functional elements underlying non-coding sequences by experimentally profiling biochemical marks across a wide array of tissues, cell lines and cell types^6–9^. An integrated encyclopedia of candidate non-coding regulatory elements, such as promoters, enhancers and silencers, has been systematically identified via multimodal molecular signatures, including DNA methylation, transcription factor (TF) binding, histone modification, chromatin accessibility and gene transcription^9^. However, the functional sequencing approaches remain fundamentally descriptive rather than predictive. These methods catalogue chromatin states based on biochemical signatures, instead of learning the regulatory code underlying native sequence, and therefore lack the generalizability to predict the function of arbitrary sequences within specific cellular contexts.

Recent advances in deep learning-based sequence-to-function models have demonstrated remarkable success in learning the relationship between genomic sequences and their corresponding regulatory functions^10,11^. Those models are capable of building predictive mapping from paired DNA sequences and experimental functional genomic data, and further uncovering mechanistic cis-regulation elements and gene regulation rules underlying genomic sequence. Models like DeepBind^12^, DeepSEA^13^, Basset^14^, SpliceAI^15^, Akita^16^ and Orca^17^ have successfully decoded diverse genomic features directly from sequence, covering a spectrum of functional profiles from transcription factors binding specificities, chromatin accessibility and Histone marks, to mRNA splicing and three-dimensional genome architecture. Furthermore, advancements like TF-MoDISco^18^, AI-TAC^19^, BPNet^20^, Nvwa^21^ and scBasset^22^ have extended the capabilities for deeper interpretability by dissecting underlying regulatory syntaxes with single-cell specificity and single-nucleotide resolution. However, current sequence-to-function models are restricted to specific genomic, functional and cellular contexts, limiting their scope to task-specific cis-regulatory elements. We still lack a comprehensive framework that learns a universal representation of genomic sequences across extensive biochemical profiles and cellular conditions, providing the foundation for integrating multi-scale understanding to construct a global encyclopedia of the entire regulatory genome.

Within the descriptive framework, functional sequencing data are systematically integrated to build a categorized vocabulary of the regulatory genome. Methods like ChromHMM^23^ and Segway^24^ define distinct chromatin states of genomic sequences based on combinatorial histone marks, allowing consortia like ENCODE and Roadmap to generate a foundational registry of candidate cis-regulatory elements across diverse cell types and tissues. Analogously, the sequence-to-function model Sei proposes the concept of sequence classes^25^. By predicting 21,907 chromatin profiles from genomic sequence alone, Sei systematically assigns genomic sequences into several distinct functional categories, and quantifies the regulatory impact of variants on sequence classes. However, the framework of Sei predominantly focuses on predicting variant effects, rather than elucidating the intrinsic characteristics and transcriptional regulatory mechanisms of genomic sequence classes. Furthermore, constrained by the CNN architecture, Sei processes only a 4-kilobase (kb) sequence window to predict the binary presence or absence for TF binding, histone marks, and chromatin accessibility at the peak center. Consequently, the restricted genomic receptive field, combined with insufficient coverage of cell types and functional profiles, limits the internal representation required for a comprehensive regulatory encyclopedia.

Conversely, sequentially predictive models such as Basenji^26^ and Enformer^27^ are designed to predict quantitative regulatory activity across broader genomic windows. Enformer leverages transformer-based architecture to expand the effective receptive field up to 100 kb, and predict 5,313 quantitative regulatory tracks across extensive cellular contexts. Most recently, AlphaGenome^28^ scales the genomic receptive field to 1 megabase while maintaining single-base pair resolution, which also captures a comprehensive array of functional contexts, from 3D chromatin contact maps to RNA splicing dynamics, effectively unifying diverse regulatory layers into a single predictive model. Crucially, sequential models process the entire genomic sequence to predict diverse functional profiles, which provide the optimal foundational architectures for constructing a global regulatory encyclopedia for the entire regulatory genome across diverse functional and cellular contexts.

Furthermore, a primary objective of learning the comprehensive representations of functional genomic sequences is to decipher the cis-regulation mechanisms of gene expression, specifically to understand how regulatory sequences orchestrate distinct gene expression programs across different cell types^29–31^. Sequence-to-function models have also demonstrated the capacity to predict gene expression directly from genomic sequences. For instance, ExPecto^32^ established an ab initio deep learning framework to predict tissue-specific gene expression and prioritize causal variants in disease loci. Enformer and AlphaGenome substantially improved accuracy of transcription signal and variant effect prediction by integrating long-range genomic information^27,28^. However, the inherent black-box nature of deep learning-based sequence-to-function models limits our ability to elucidate transcription regulation rules, especially for the interaction grammar between functional cis-regulatory elements.

Here, we address these challenges by introducing HERMES (Hierarchical Encoding of Regulatory Mechanisms and Expression Syntax), a foundational framework trained on a massive compendium of 137,127 functional genomics profiles, spanning DNA methylation, TF binding, polymerase binding, histone marks, chromatin accessibility and RNA expression across diverse tissues, cell lines and cell types, to construct a unified, quantitative map of the entire genome. HERMES established a context-specific regulatory vocabulary by comprising 40 distinct sequence classes through unsupervised clustering of learned functional representations, and further defined fine-grained subclusters for promoter and enhancer sequence class. By interrogating this vocabulary, we deciphered the sequence determinants of a diverse repertoire of highly tissue-specific enhancer classes, and identified a stable promoter class governing housekeeping genes driven by ETS, YY1 and CCAAT motifs. Crucially, we leveraged the high-dimensional sequence embedding to transcription regulation syntax by predicting gene expression across 3087 tissue or cell-type-specific conditions. We validated the regulatory perturbation prediction using CRISPRi perturbation datasets, and further inspected the regulatory architecture of each gene by quantifying its sensitivity to non-coding enhancer perturbations. Ultimately, we distilled the deep learning-based sequence-to-function HERMES model into a biologically interpretable transcriptional regulation formula, providing an explicit quantitative rulebook that reveals the underlying mechanism of promoter-enhancer interactions. In total, we propose a hierarchical encoding framework to decipher the comprehensive regulatory mechanisms underlying native genome sequences, by progressively defining a fundamental sequence vocabulary, parsing the gene expression syntax into a quantitative rulebook for complex regulatory genomes.

## Results

### Build a foundational sequence-to-function model

To comprehensively learn the universal representation of the entire genomic sequences across extensive functional and cellular contexts, we developed a foundational deep-learning-based sequence-to-function model named HERMES, which was designed to predict a broad spectrum of functional genomics data and encompass the long-range context information from genomic sequences.

In order to decode the diverse regulatory function of the genome sequences, HERMES model was trained to predict a total of 137,127 distinct functional genomics sequencing profiles across comprehensive functional and cellular contexts. Specifically, the dataset was stratified into 106,663 peak-based and 30,464 signal-based assays. The peak-based data provided a multi-faceted view of the regulatory landscape, including 30,408 histone marks, 27,281 transcription factor (TF) binding, 23,131 chromatin accessibility, 19,022 DNA methylation, 3,398 polymerase binding and 2,322 chromatin accessibility profiles mainly from ChipAtlas^33^ database and RoadMap project. Quantifying the intensity of regulatory activity, the signal-based data was composed of 8,802 histone marks, 8,382 TF binding and 2,033 chromatin accessibility profiles from Cistrome^34^ database (19,217 in total), 2,530 TF binding, 2,428 histone marks, 1,420 chromatin accessibility and other regulatory profiles from Encode project (7,273 in total), 2,327 RNA Expression and 1,423 CAGE profiles mainly from Encode and Fantom^35^ project (Fig. 1a). A key strength of the large-scale dataset lies in its structured heterogeneity, characterized by a broad range of distinct biochemistry profiles and biological sources. Specifically, the dataset integrates biochemical assays spanning DNA methylation, TF binding, polymerase binding, histone marks, chromatin accessibility and RNA expression. Furthermore, it encompasses a wide spectrum of biological contexts, ranging from primary tissues, stem cells, organoids to diverse cell lines (Fig. 1b). The heterogeneity of the curated training data provides a robust foundation for learning a generalizable model of regulatory genomic function across diverse biological contexts.

**Figure 1.**
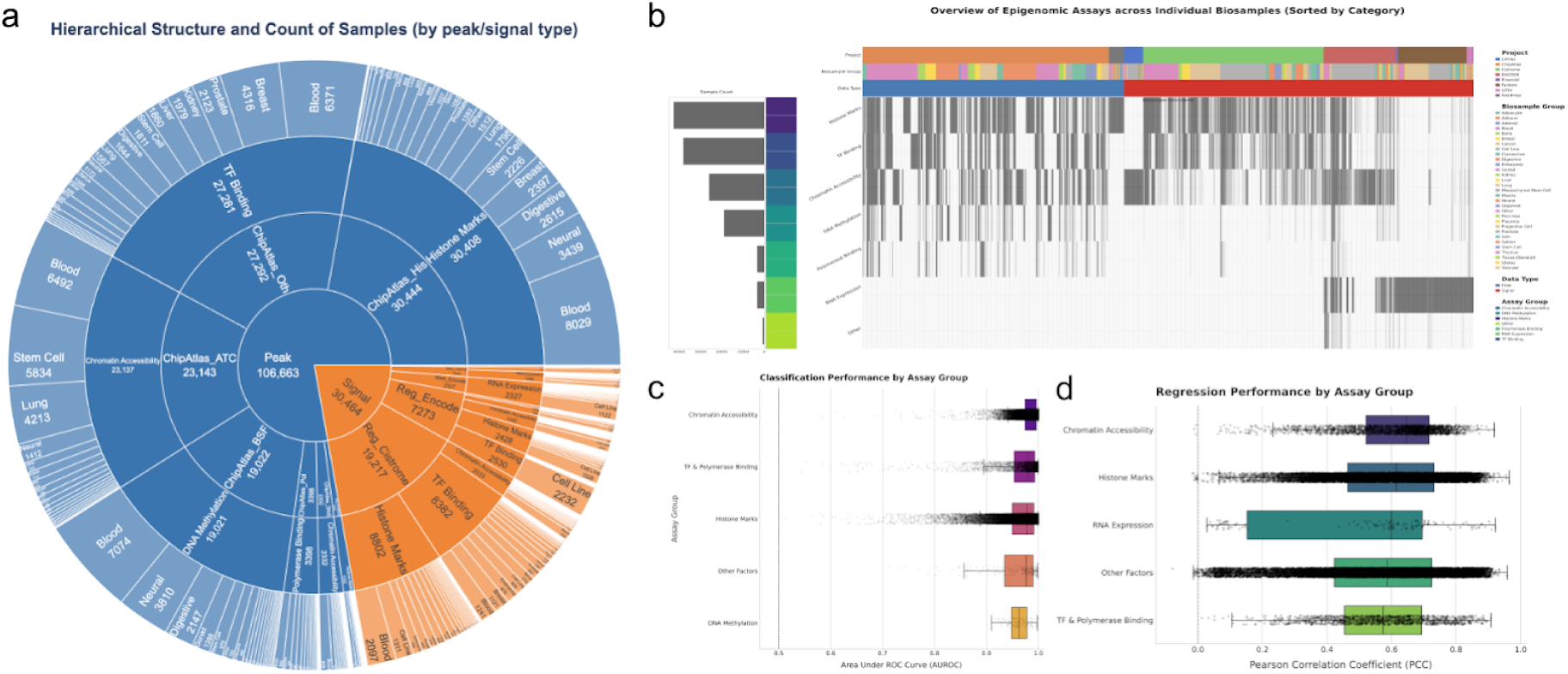
**Overview of the HERMES dataset curation and model performance.** a, Hierarchical structure and sample counts of the training dataset. The HERMES model was trained on a comprehensive set of 137,127 functional genomics profiles, stratified into 106,663 peak-based assays comprising histone marks, TF binding, chromatin accessibility and DNA methylation and 30,464 signal-based assays including regulatory profiles, RNA expression and CAGE profiles. b, Visualization of data distribution across 859 samples (columns) and 40 assays (rows), ordered by the number of experiments (indicated in parentheses) and colored by sample metadata. c, Classification performance by assay group. Discriminative evaluation of peak-based tasks measured by Area Under the ROC Curve (AUROC) and Area Under the Precision-Recall Curve (AUPRC). The model demonstrated high accuracy with an average AUROC of 0.972, showing excellent discrimination across TF binding, chromatin accessibility and histone mark profiles. d, Regression performance by assay group. Quantitative evaluation of signal-based tasks using Pearson (PCC) and Spearman (SCC) correlation coefficients. The model achieved a robust average PCC of 0.57 across all signal-based tasks, with notably high performance in RNA expression and chromatin accessibility prediction.

To encode the genomic sequence, HERMES utilized the Enformer-like architecture with the receptive field of 153.6kb to integrate information across distal elements. Through ablation study, we updated the convolution size for better feature capturing, and adopted a self-gated attention module to control the information fusion (SFig. 1a). We also designed related auxiliary tasks by predicting conservation scores of PhaseCon and PhyloP from genomic sequence, in order to increase the representative learning ability of the HERMES model with evolutionary information (SFig. 1b).

After training, HERMES model achieved accurate predictive performance among all the functional profiles on the test set of held-out chromosomes. Across all peak-based tasks, the model received an average area under the receiver operating characteristic (AUROC) of 0.98 (Fig. 1c) and average area under the precision-recall curve (AUPRC) of 0.41 (SFig. 2c). For all signal-based tasks, the average Pearson correlation coefficient (PCC) was 0.57 (Fig. 1d) and Spearman correlation coefficient (SCC) of 0.51 (SFig. 2d). Furthermore, HERMES model demonstrated remarkable versatility across a diverse spectrum of prediction tasks, with predictive performance consistently maintaining high accuracy across different functional assay groups. For peak-based classification, HERMES model achieved robust performance for Histone Marks (mean AUROC=0.98) , Chromatin Accessibility (mean AUROC=0.99), TF Binding and Polymerase Binding (mean AUROC=0.99), DNA methylation (mean AUROC=0.96) and Other Factors (mean AUROC=0.98). In terms of quantitative signal prediction, the model also shows robust fidelity across tasks. The highest correlations were found for RNA Expression (mean PCC=0.50), Chromatin Accessibility (mean PCC=0.62), TF and Polymerase Binding (mean PCC=0.58) and Histone Marks (mean PCC=0.61). By visualizing representative genomic signal tracks at held-out genomic regions, we validated the high concordance between predicted and observed genomic profiles (SFig. 2e).

**Figure 2.**
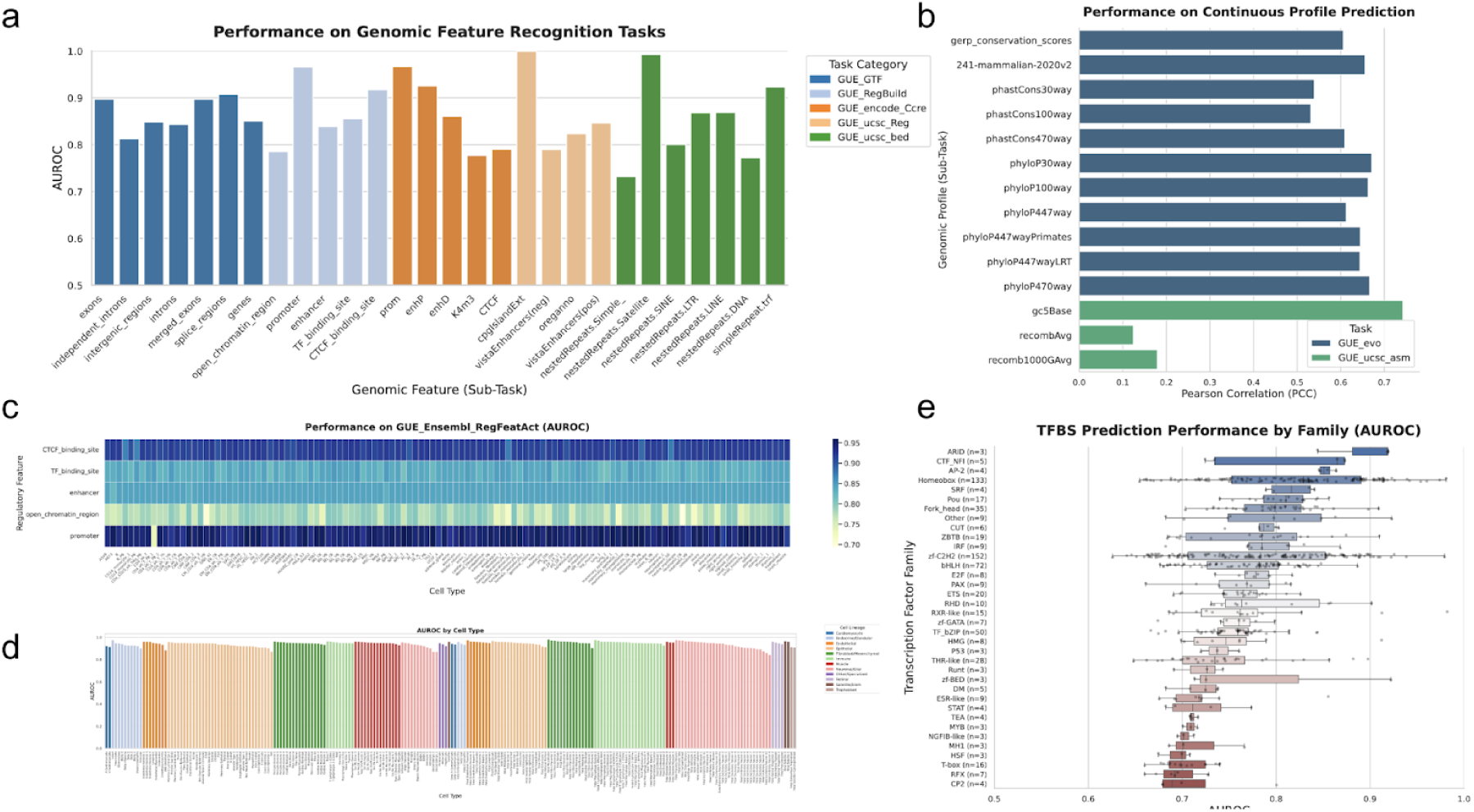
**Evaluation of predictive performance across annotated genomic features and regulatory contexts.** a, Classification performance on genomic features. Bar plot showing the Area Under the Receiver Operating Characteristic (AUROC) for peak-based prediction tasks, stratified by different categories of genomic features. b, Quantitative signal prediction accuracy. Pearson correlation coefficients (PCC) assessing the agreement between predicted and observed signal intensities across conserved genomic signal regions and genomic features. c, Performance on Ensembl regulatory features. Heatmap displays AUROC scores for the prediction of regulatory feature activity (RegFeatAct), categorizing performance across specific element types and cell types defined by Ensembl annotations. d, Lineage-specific performance on cis-regulatory elements. AUROC scores for predicting candidate cis-regulatory elements (cCREs) from the Catlas database, stratified by cell lineage. This illustrates the model’s capability to capture tissue-specific regulatory landscapes. e, Transcription factor binding prediction by family. Distribution of AUROC scores for Transcription Factor Binding Sites (TFBS), grouped by transcription factor families, highlighting the model’s robustness across diverse DNA-binding mechanisms.

In addition, we further evaluated the consistency of HERMES’ predictive performance across different genomic regions, by calculating the average accuracy metric across functional profiles for all the test sequences on held-out chromosomes. The performance remained robust and consistent along the chromosome arms across all the signal and peak tasks. Notably, we observed a decrease in performance specifically within the centromeric regions of the whole chromosome. This localized degradation was likely attributable to the highly repetitive nature of centromeric DNA (SFig. 3a-c), which poses known challenges for sequence alignment and data quality, thereby impacting predictive accuracy^36^.

**Figure 3.**
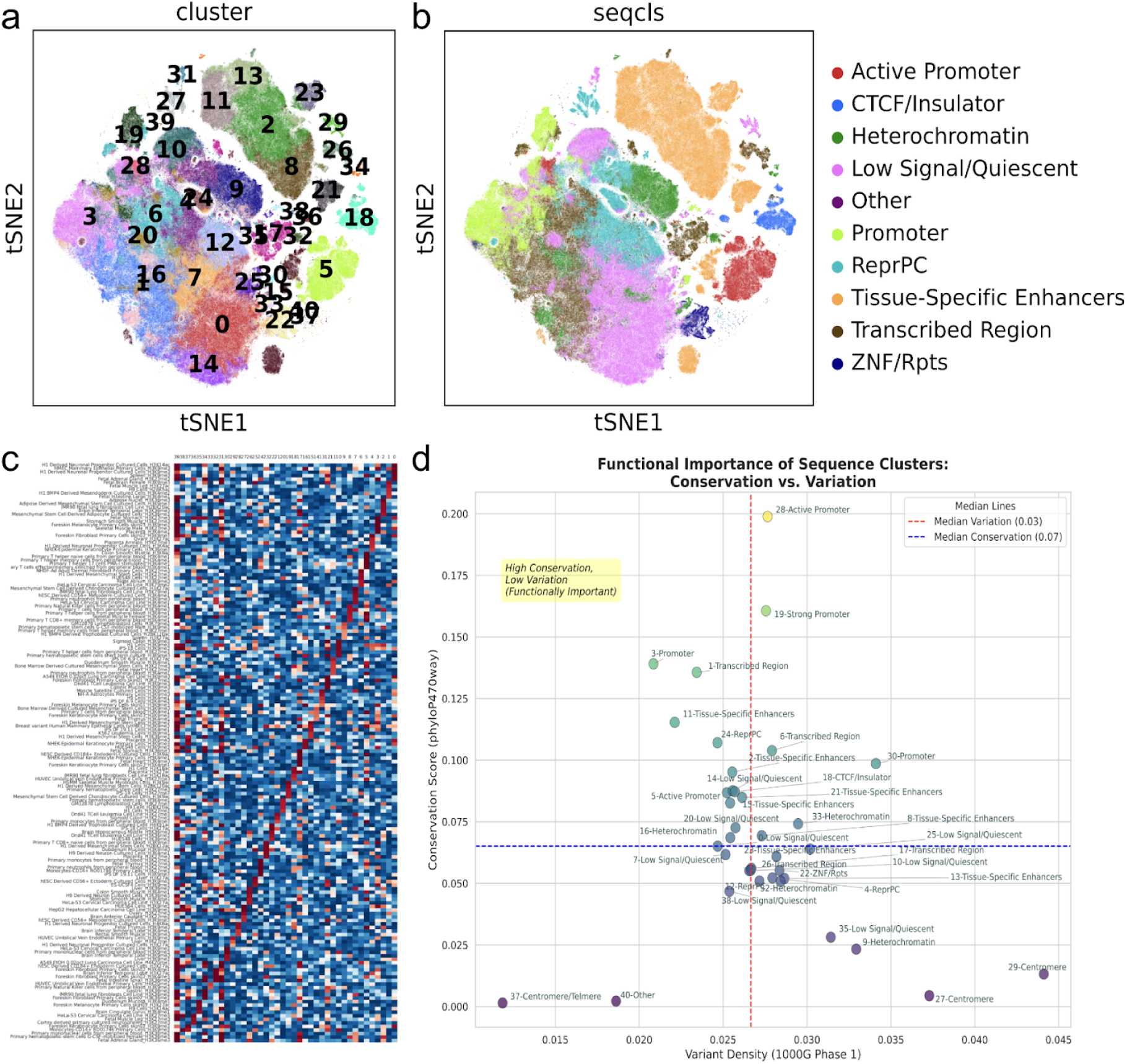
Unsupervised discovery of functional sequence classes from HERMES embeddings. a, Global clustering of genomic sequences. The HERMES model embeddings for 128-bp sequence tiling the entire genome were projected and clustered, revealing a highly structured regulatory landscape consisting of 40 distinct sequence classes. b, Functional annotation of sequence classes. The identified clusters were annotated based on their dominant biological features and regulatory roles. c, Characterization of regulatory signatures. Heatmap displaying the enrichment of each sequence class within the Roadmap Epigenomics biochemical profiles. This analysis reveals the molecular basis of the sequence classes, with each cluster displaying a unique signature of regulatory marks corresponding to distinct tissue states. d, Evolutionary constraints and functional importance. Scatter plot comparing the evolutionary conservation score against genetic variation density for each sequence class. The distribution highlights classes under strong purifying selection (high conservation, low variation), indicating their functional importance in the genome.

Beyond accurately predicting individual profiles, HERMES predictions also preserved the intrinsic structure between correlated tasks. By visualizing the task representations using the hidden embedding weights of the output layer, we found that biologically related assays could spontaneously group into the same clusters (SFig. 4b,e), while also capturing fine-grained tissue and cell-type specificities among closely related samples (SFig. 4c,f).

**Figure 4.**
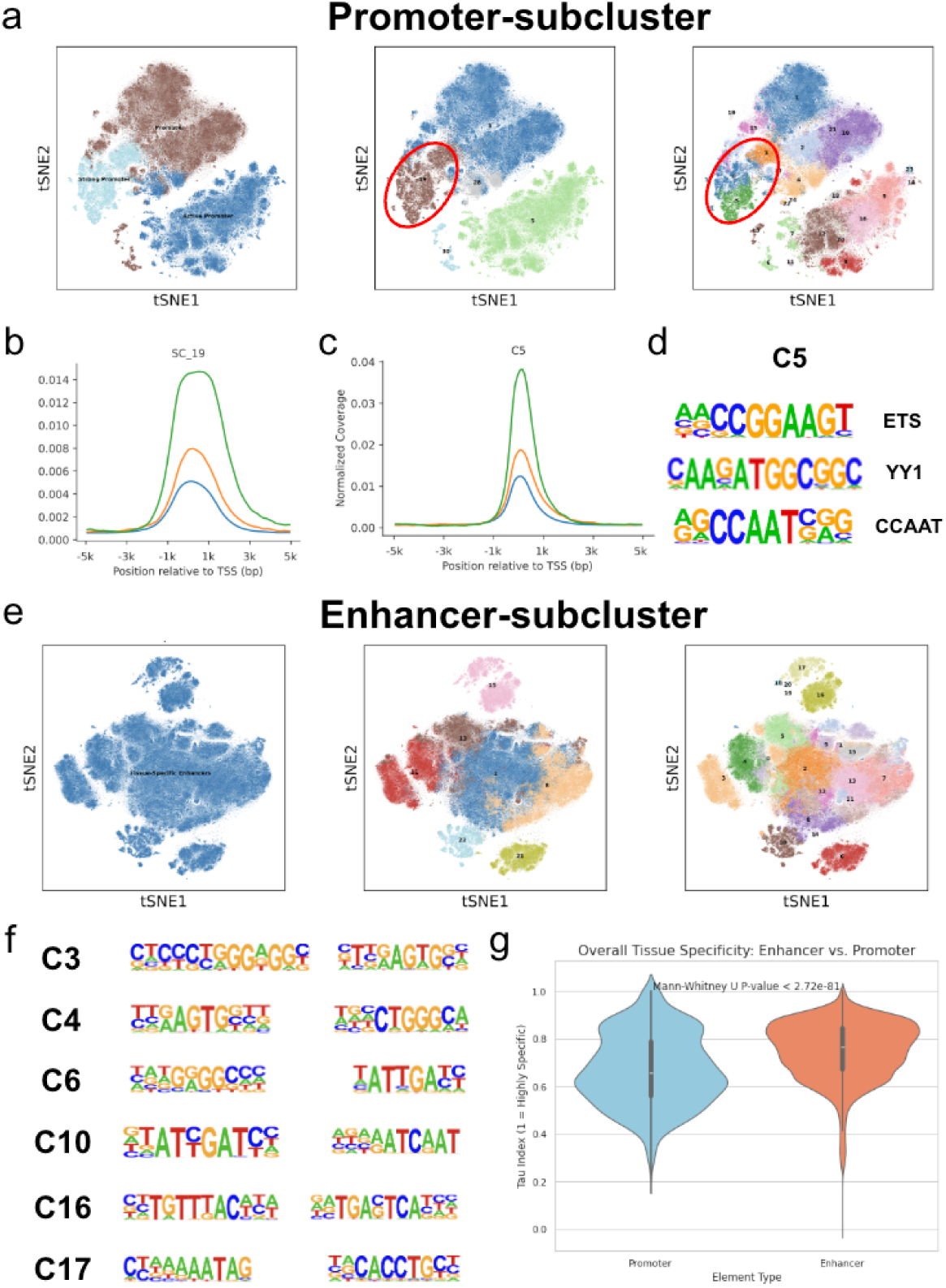
**Hierarchical clustering of Promoter and Enhancer Sequence Class identified fine-grained subclusters and underlying motifs.** a, The t-SNE visualizations of Promoter sequence class and subclusters. The left and middle panel shows the initial classification of Promoter sequence class. The right panel reveals the refined sub-clusters identified using the Leiden algorithm. b, Profile plots visualizing the spatial distribution and normalized enrichment coverage of the sequence classes C19 within a ±5 kb window centered on the Transcription Start Sites (TSS). c, Profile plots visualizing the spatial distribution of the promoter subcluster C5 within a ±5 kb window centered on the TSS. d, Motif enrichment for each Promoter subcluster C5. The sequence logos provide a visual representation of the core enriched motifs. e, The t-SNE visualizations of Enhancer sequence class and subclusters. The left and middle panel shows the initial classification of Enhancer sequence class. The right panel reveals the refined sub-clusters identified using the Leiden algorithm. f, Motif enrichment for each Enhancer subcluster. The sequence logos displayed next to several cluster IDs provide a visual representation of the core enriched motifs. g, Tau index comparison of Promoter sequence classes and Enhancer sequence classes based on the chromatin accessibility among cell types.

In summary, HERMES represents the most comprehensive sequence-to-function framework established to date, which predicts a total of 137,127 distinct functional profiles covering extensive functional and cellular contexts. The rigorous evaluation confirms that the model has successfully learned a comprehensive mapping from genomic sequences to complex functions of the regulatory genome, effectively capturing the underlying sequence patterns and intrinsic co-variation of different regulatory landscapes.

### Sequence representations understand diverse genomic features

To rigorously assess the semantic richness and generalizability of the sequence representations learned by HERMES, we established a systematic framework to evaluate the model’s genome understanding capacity. To this end, we leveraged the sequence embeddings from pre-trained HERMES to large-scale genomic annotations as a comprehensive suite of downstream supervised tasks. The evaluation suite was curated to encompass the biological complexity of the genome, spanning fundamental genomic properties, gene entities, regulatory elements, and cross-species evolutionary constraints. Crucially, these tasks ranged from the recognition of static sequence features to the prediction of cell-type-specific activities, and further evaluated the extent to which sequence embeddings capture the fundamental information content of the human genome.

First, we assessed the model’s capacity to recognize basic genomic features from sequence alone (Fig. 2a; SFig. 5a). HERMES demonstrated high accuracy across diverse categories, achieving a median AUROC of 0.85 for tasks related to gene structure. For identifying core regulatory elements, AUROC predictive performance was 0.95 for promoter, 0.92 for enhancer and 0.99 for CPG island from RegBuild, Encode_cCRE and UCSC Regulatory database. The model also effectively learned to distinguish various repeat elements, such as SINE (AUROC = 0.99) and LINE (AUROC = 0.88), showcasing a fundamental understanding of genomic architecture. We next tested the model’s ability to predict the evolutionary properties of the genome (Fig. 2b; SFig. 5b). HERMES accurately predicted cross-species conservation scores, achieving high Pearson correlations for both phyloP (mean PCC = 0.68) and phastCons (mean PCC = 0.55) profiles. However, prediction accuracy was notably lower for cross-individual recombination rates (PCC = 0.18), demonstrating the limited predictive power of sequence-to-function models to capture the distinct sequence determinants driving population-level recombination dynamics.

**Figure 5.**
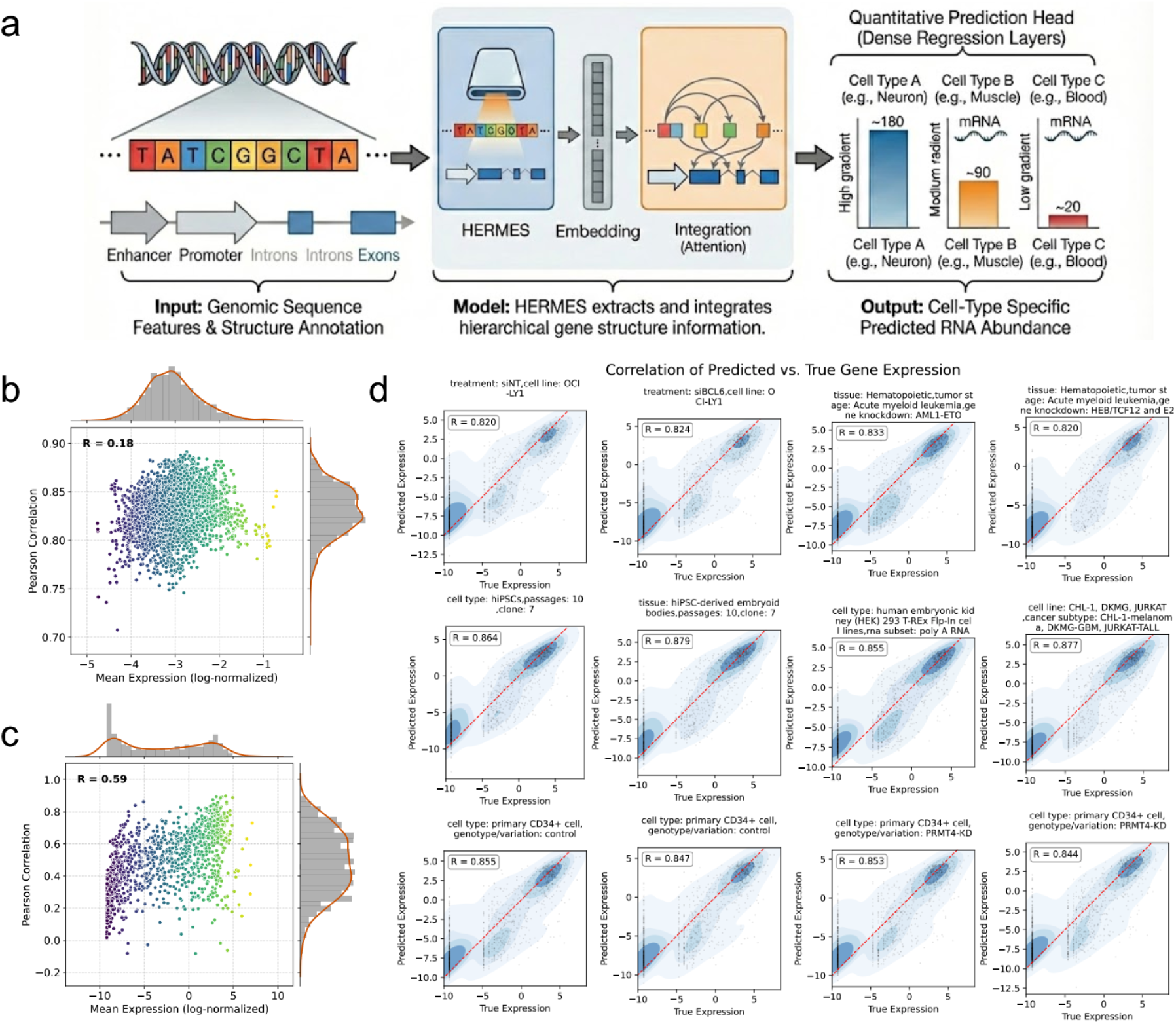
**Architecture and quantitative evaluation of sequence-based gene expression prediction.** a, Schematic overview of the model architecture adapted for quantitative gene expression prediction. The diagram illustrates how the model integrates genomic sequence features to predict RNA abundance levels. b, Per-task performance consistency across biosamples. Scatter plot displaying the per-task predictive accuracy (Pearson correlation coefficient, PCC) for each cell type/biosample against its mean expression level. This analysis assesses whether model performance is robust across samples with varying sequencing depths or library sizes. c, Per-gene performance dependence on gene expression levels. Scatter plot correlating the per-gene predictive accuracy (Pearson correlation coefficient, PCC) with the mean expression level of that gene. This evaluates the model’s ability to predict expression across a dynamic range of expressed genes. d, Validation on holdout chromosome 8 across representative tissues. Scatter plots compare the sequence-based predicted expression (x-axis) against experimentally measured RNA-seq values (y-axis) for 990 genes on the holdout chromosome 8. Results are shown for six example tissues, with Pearson correlation coefficients (PCC) indicating a high concordance between predicted and observed expression profiles. *The illustration was drawn by Gemini Nano Banana*.

A more challenging test is the prediction of cellular condition-specific regulatory elements. We evaluated HERMES on the RegFeatAct task, which requires predicting the activity of promoters, enhancers and CTCF sites across a wide panel of cell types. The model showed exceptional and highly uniform performance, with median AUROCs > 0.90 for all regulatory element types (Fig. 2c; SFig. 5c). This high accuracy was consistent across the entire spectrum of cell types (Fig. 2d; SFig. 5d), indicating that HERMES has learned a generalizable, cell-type-aware regulatory code.

Finally, given the critical role of transcription factors (TFs) in defining cell state, we benchmarked the model’s ability to predict binding sites, stratified by TF family (Fig. 2e; SFig. 5e). HERMES demonstrated strong performance across the board, with particularly high accuracy for families with well-defined structural motifs, such as Homeodomain (median AUROC = 0.84), ForkHead (median AUROC > 0.80) and ARID (median AUROC > 0.92). This robust performance, even for more complex families, confirmed a deep understanding of TF binding syntax.

Collectively, our comprehensive evaluation demonstrates that HERMES exhibits exceptional robustness and versatility across a broad spectrum of genomic understanding scenarios, and the model embedding has successfully distilled a rich and biologically meaningful representation of genome sequences. The foundational representation learning ability of HERMES motivates us to leverage the high-fidelity sequence embedding to uncover the intrinsic organization of the genome.

### Define comprehensive sequence vocabulary of the entire genome

The interpretation of the human genome hinges upon understanding the sequence-intrinsic representations of underlying regulatory activity. The HERMES model was designed to predict an extensive collection of functional profiles directly from DNA sequences, yielding a context-specific functional vocabulary of the human genome. Using the foundational model, we further constructed a comprehensive catalogue that systematically annotates these fundamental functional elements, providing a map of their activity across a wide spectrum of biochemistry assays and tissue environments.

To systematically categorize the functional landscape of the human genome, we first leveraged the pretrained HERMES model to generate high-dimensional sequence embeddings across the entire genome. In total, we extracted approximately 30 million sequence embeddings with 3052 hidden dimensions that uniformly tile the entire genome at a 128bp resolution. These context-aware representations integrated comprehensive information from 137,127 distinct functional genomics profiles, while capturing long-range dependencies across a broad 153.6kb receptive field. T-SNE^37^ visualization of the sequence representations revealed a highly structured global organization of regulatory activities encoded in the genomic sequences (Fig. 3a,b). We further applied community clustering to partitioning the genome into 40 distinct sequence classes (Fig. 3a, Methods) and annotated each cluster based on its specific enrichment for histone marks and transcription factor binding profiles (Fig. 3b, Methods). Collectively, these sequence classes provided a near-complete annotation of the human genome, covering >99.6% of the human genome.

To resolve the functional identity of the sequence classes, we enriched the chromatin marks for each sequence cluster. Heatmap elucidated that different branches of sequence classes are enriched in distinct chromatin modifications (Fig. 3c). For example, Clusters C3, C5, C19, C28, C30 mapping to the Promoter group showed broad enrichment for active promoter marks (e.g., H3K4me3, H3K9ac, H3K27ac). The large Tissue-Specific Enhancer clusters (C2, C8, C11, C13, C15, C21, C23) were characterized by highly cell-type-specific enrichment of H3K4me1, H3K27ac marks. The model precisely distinguished between different repressive states. Heterochromatin clusters (C9, C16, C32, C33, C34) were defined by H3K9me3 enrichment. In contrast, Polycomb-repressed (ReprPC) clusters (C4, C12, C24) were clearly marked by H3K27me3. Other distinct functional elements, such as Transcribed Regions C1, C6, C17, C26 were also successfully identified and separated based on H3K36me3.

Furthermore, we systematically validated the annotation of sequence class by enrichment for a wide range of independent genomic annotations, including gene-centric features, curated regulatory regions and genomic properties (SFig. 6). The clusters partitioned cleanly based on known genomic and regulatory features. For example, Clusters C5, C19 and C28 were all enriched for encodeCombined.promoter and cpgIslandExt, linking them to the promoter-associated and CpG islands-associated regulatory functions. Clusters C13, C15, C21 were highly enriched for experimentally validated encodeCombined.enhancer and vistaEnhancers, confirming their identity as a class of active enhancers. Genomic annotations further help to characterize the biological identity of each sequence class. For instance, Clusters C18 and C36 showed specific and strong enrichment for encodeCombined.CTCF sites, identifying them as insulator or CTCF-associated classes. Clusters C27, C29, C31, C34, C36 were associated with Centromere and Telomere. Cluster C22 showed unique enrichment for nestedRepeats.

**Figure 6.**
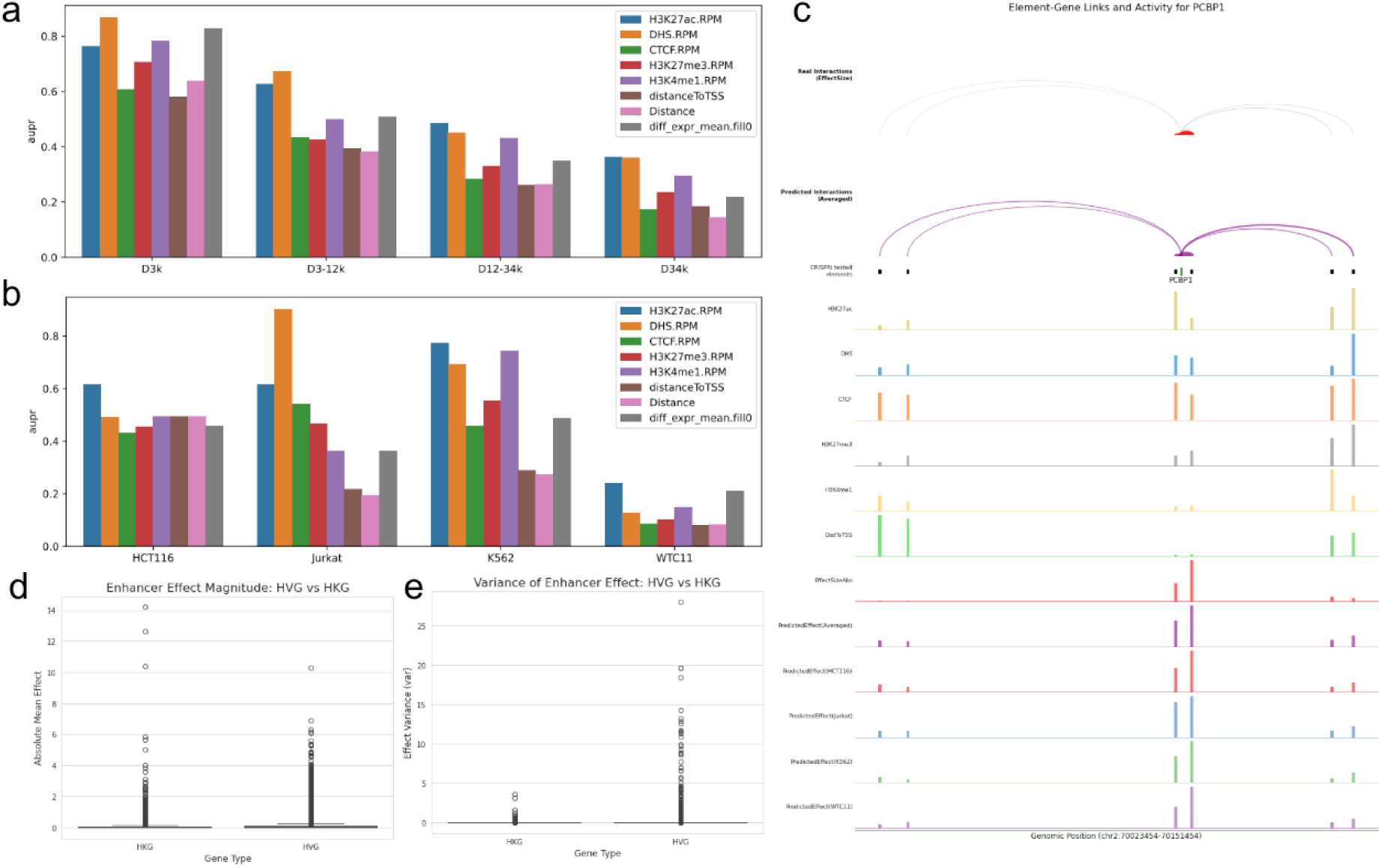
**Validation of enhancer-gene regulatory predictions and characterization of gene-type specific regulatory architectures.** a, Genomic distance stratified benchmarking on K562 CRISPRi datasets. Evaluation of enhancer-gene pair classification accuracy (CRISPRi-validated positives versus non-validated candidates) measured by the Area Under the Precision-Recall Curve (AUPRC). Performance is assessed using individual chromatin features (H3K27ac, DHS, CTCF, H3K27me3, H3K4me1 and model prediction) and stratified by the relative distance between the candidate enhancer and the target gene TSS, highlighting the predictive performance at varying distances. b, Cross-cell-type benchmarking on CRISPRi datasets. AUPRC scores assess the model’s ability to identify functional enhancer-gene pairs across four distinct cell lines (HCT116, Jurkat, K562 and WTC11). This demonstrates the robustness of the predictions across diverse cellular contexts. c, Visualization of the PCBP1 regulatory landscape. Example arc-plot anchored at the PCBP1 gene, illustrating the high-confidence long-range regulatory interactions (loops) predicted by the model. H3K27ac, DHS, CTCF, H3K27me3, H3K4me1 chromatin features, and cell-type specific CRISPRi effects were plotted according to genomic positions. All the 3084 cell type-specific enhancer effect size predictions were visualized in heatmap according to the same genomic positions. d, Comparison of enhancer effect magnitude for HKGs and HVGs. Box plots displaying the distribution of predicted enhancer effect sizes (regulatory impact) for Housekeeping Genes (HKG) versus Highly Variable Genes (HVG). e, Analysis of regulatory variance for HKGs and HVGs. Comparison of the variance in enhancer effects for HKGs and HVGs, highlighting the differential regulatory plasticity and dynamic range between constitutive and variable gene classes.

Taken together, the t-SNE visualization of the genomic sequence classes showed a clear organization with known biological relationships. For instance, sequence classes with weak or no regulatory activity (Low Signal/Quiescent) form the central core of the whole genome. Core regulatory elements, such as Tissue-Specific Enhancers and Promoters sequence classes, formed distinct yet related branches, highlighting the unique sequence features that define their functions. Similarly, repressive sequence classes, including CTCF/Insulator, Heterochromatin and Repressed Polycomb (ReprPC), were located adjacent to one another, consistent with their cooperative roles in gene silencing.

Finally, we systematically assessed the evolutionary conservation and functional tolerance of the sequence class. Specifically, we analyzed the evolutionary properties for each sequence class by comparing the cross-species evolutionary conservation against their tolerance for genetic variation within the human population. As expected, indispensable regulatory elements like Active Promoters (cluster C28) and Strong Promoters (cluster C19) were characterized by high conservation and low variant density, indicating strong negative selection and a critical, constrained function. In contrast, classes associated with structurally dynamic regions, such as Centromeres (cluster C29) and ZNF/Repeats (cluster C22), exhibited very low conservation and high variant density. Interestingly, Tissue-Specific Enhancers (e.g., clusters C8, C18) occupied a distinct space of high conservation but also higher-than-median variation, potentially reflecting their role in species-specific adaptations (Fig. 3d; SFig. 7).

**Figure 7.**
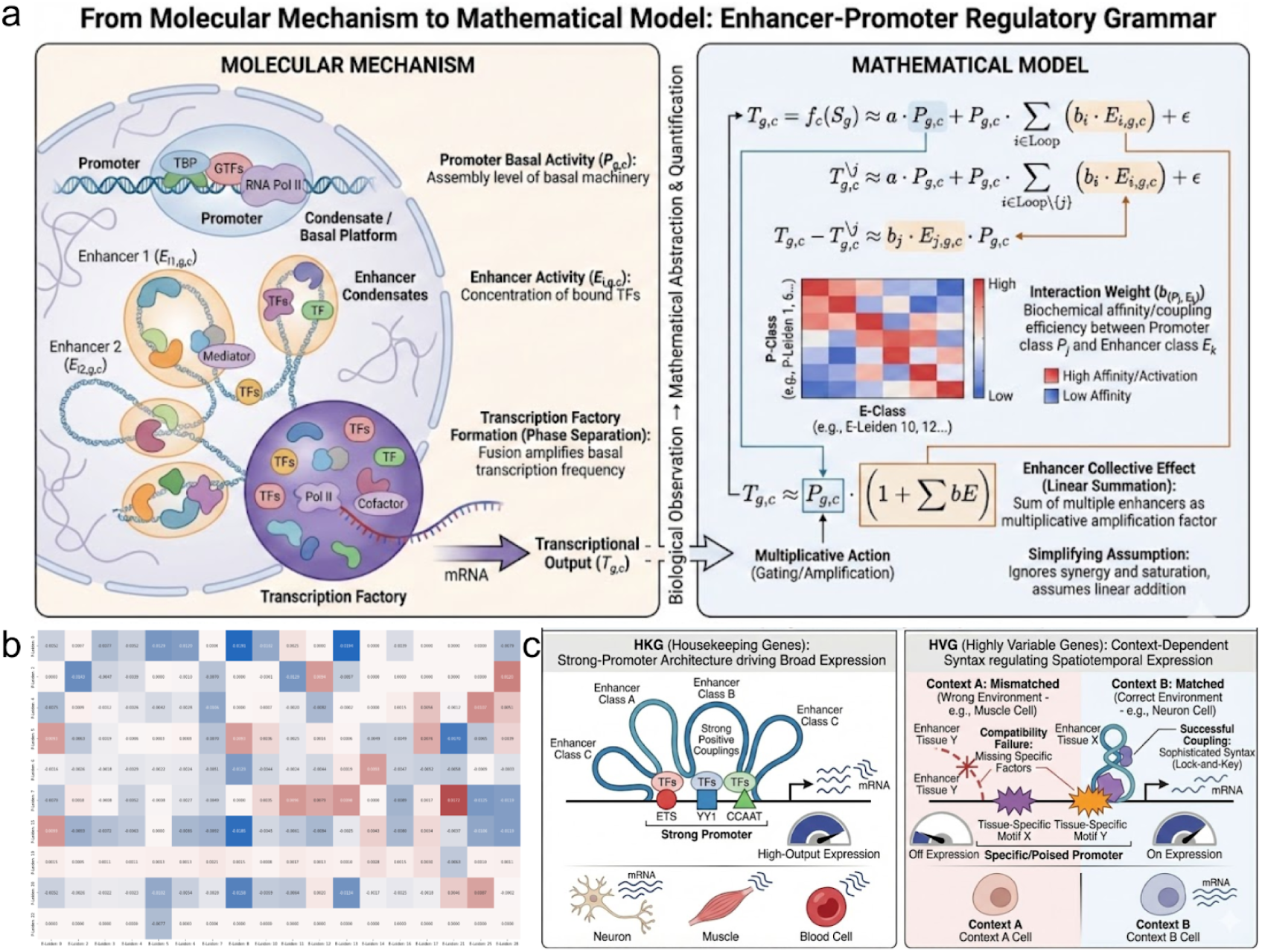
**Linear formula and Compatibility rules of gene expression.** a, From molecular mechanisms to a mathematical formula of E-P regulatory grammar. Left: Schematic representation of the molecular machinery in the nucleus. Right: Mathematical abstraction of the biological process into a full-interaction linear model. b, Pairwise interaction landscape between promoter and enhancer sequence classes. Heatmap displaying the estimated coupling efficiency (interaction weights) between distinct sequence classes of promoters (rows) and enhancers (columns). c, A unified compatibility rule of transcriptional regulation. Left: Housekeeping genes (HKGs) utilize a promoter architecture enriched with constitutive motifs (ETS, YY1 and CCAAT). These promoters exhibit high intrinsic activity and universal compatibility, forming strong positive couplings with diverse enhancer classes to maintain stable, high-output expression across various cellular contexts. Right: Highly variable genes (HVGs) rely on a context-dependent syntax. Their promoters and enhancers are characterized by tissue-specific motifs that require matching transcription factors. In mismatched contexts (Top), the absence of specific factors leads to compatibility failure and gene silencing. In matched contexts (Bottom), the presence of cognate factors enables successful E-P coupling, orchestrating precise spatiotemporal expression. *The illustration was drawn by Gemini Nano Banana*.

By harnessing the foundational representation capability of HERMES, we have constructed a comprehensive vocabulary that systematically annotates fundamental functional elements of the entire genome, providing a global catalogues of sequence classes across a wide spectrum of biochemical assays and tissue environments. The identified sequence class, including promoters, enhancers, insulators, transcribed regions, repressive elements and repeat families, constitutes a vocabulary of regulatory elements that orchestrate sequence to their corresponding biochemical functions. Furthermore, the coherence of sequence classes underscores the interplay of evolutionary conservation and functional tolerance, highlighting the selective pressures that have shaped the genome’s functional landscape.

## A promoter sequence class specific to House-Keeping Genes

To investigate the heterogeneity within the broadly defined Promoter sequence class, we performed a secondary clustering analysis on all sequences previously assigned to the Promoter category. This approach aimed to resolve broad Promoter categories into more fine-grained, functionally coherent modules that correspond to specific cellular contexts and developmental programs.

Sub-clustering revealed distinct functional archetypes of promoters based on their underlying sequence features, suggesting significant underlying diversity (Fig. 4a). We further clustered Promoter class into 25 subclusters, each of which likely represents a distinct functional archetype. As expected, subclass C3, C4, C7 and C8 showed consistent enrichment for the active promoter mark H3K4me3 across all analyzed cell types, confirming its function as broadly active promoters. In contrast, subclass C5, C6, C8, C9, C17 and C19 exhibited strong cell-type specificity. They were enriched for active enhancer marks (H3K4me1 and H3K27ac) in specific cell types, such as brain, muscle, monocytes and stem cells. Notably, regulatory regions of subclass C1, C2 showed strong enrichment for the repressive mark H3K27me3 in inactive cell types, validating the mechanism of Polycomb-mediated cell-specific gene silencing.

To uncover the molecular basis for the sub-classification, we performed a motif enrichment analysis for each newly identified promoter sub-cluster. Motif Enrichment also identified functionally distinct TFBS underlying fine-grained promoters (SFig. 8a,b). For example, we annotated several key sub-clusters based on their dominant motifs. Subclass C0 was strongly characterized by the presence of the TATA-Box, the binding site for the TATA-binding protein (TBP), defining it as the canonical TATA-box promoter class. Subclass C3 was enriched for motifs recognized by Sp1(Zf) transcription factors, a common feature of GC-rich promoters for housekeeping genes. Subclass C5 displayed a combination of motifs for ETS, YY1 and CCAAT factors, representing another well-established class of core and proximal promoter elements. This granular analysis demonstrated that the broad Promoter class is not a monolithic entity, but rather a composite of diverse regulatory architectures.

With the anchor of Transcription Start Sites (TSSs), we next sought to validate their functional relevance in the context of gene regulation. We analyzed the enrichment patterns of sequence class in TSS flanking regions, and further compared the regulatory sequence between expressed Housekeeping Genes (HKGs) and Highly Variable Genes (HVGs). Our analysis of sequence class enrichment around TSSs confirmed that the model has learned biologically meaningful features (Fig. 4b; SFig. 9a). As expected, we observed a strong and localized enrichment of promoter-associated classes (C5, C19, C24) in the immediate vicinity of TSSs, consistent with established biological knowledge. Among all the sequence classes, C19 emerged as the class with the highest enrichment level, identifying it as a key component of core promoters.

We further dissected the enrichment of promoter related sub-clusters to reveal fine-grained distinct functional archetypes (Fig. 4c; SFig. 9b). Among the sub-clusters, C0 and C5 contributed most to the Promoter activity of HKGs, representing the sequence basis of C19 core promoter elements. However, they were architecturally and compositionally divergent. C5 exhibited a focused, unimodal enrichment peak at the core TSS, and was accordingly enriched for ETS, YY1 and CCAAT factor motifs. In direct contrast, C0 displayed a bimodal signal flanking the TSS, a classic signature of TATA-containing promoters, and was correspondingly enriched for TATA-box factor motifs.

To further validate the regulatory mechanisms underlying housekeeping and variably expressed genes, we compared their chromatin accessibility landscapes using single-cell ATAC-seq (scATAC) data^38^. The aggregate profile of scATAC signals were centered around the TSS for both HKGs and HVGs, indicating a consistently open chromatin region. Besides, the average signal intensity of HKG was substantially higher than HVG, suggesting a more constitutively open promoter architecture of HKG across all cell types (SFig. 9c). Next, we analyzed the distribution of accessibility signals across all promoter regions for each gene class. The distribution for HKGs was unimodal with a tight peak at a relatively high accessibility value, while the distribution for HVGs was distinctly bimodal with different states across tissues (SFig. 9d).

Through a secondary clustering analysis within the Promoter sequence class, we defined fine-grained sub-clusters corresponding to specific genomic and cellular contexts. Moreover, the comparative analysis between HKGs and HVGs indicates fundamentally different regulatory strategies, HKGs possess universal strong promoters, while the regulation of HVGs involves context-specific promoter elements. Specifically, we found a sub-class of promoters with underlying motifs of ETS, YY1 and CCAAT factors, driving the HKGs to maintain a stably open state across diverse cellular contexts.

### Enhancer sequence classes underscores cellular specificity

Following our analysis of promoters, we applied the same sub-clustering strategy to dissect the extensive heterogeneity within the Enhancer sequence class. The t-SNE projections and sub-clustering resolved the broad sequence class into 21 distinct functional sub-groups (Fig. 4e).

Characterization of Enhancer sub-clusters demonstrated that enhancer heterogeneity is primarily derived by cell-type specificity (SFig. 10a). These sub-clusters were distinguished by a dynamic chromatin signature, marked by the hallmark co-enrichment of H3K4me1 and H3K27ac specific to distinct tissues and cell types. For instance, subclass C4 showed strong, specific enrichment for H3K4me1 and H3K27ac in Brain and Foreskin Keratinocyte samples. Subclass C6 was similarly activated in Foreskin Fibroblasts. Subclass C8 represents a liver-specific (HepG2) active enhancer marked by H3K4me1 and H3K27ac. Notably, the model also successfully isolated poised or Polycomb-repressed enhancer archetypes, which are critical for developmental regulation and are typically marked by H3K27me3. Subclass C18 and C20 both showed strong and specific enrichment for the repressive mark H3K27me3 in progenitor and stem cell lineages. Sub-clustering further identified classes representing a primed state, such as Subclass C10, which showed broad enrichment for H3K4me1 but lacked both the active H3K27ac and repressive H3K27me3 marks.

To define the functional identity of these enhancer sub-clusters, we also performed a comprehensive motif enrichment analysis. Motif enrichment revealed that the sub-clusters are strongly defined by combinations of transcription factor binding motifs known to drive cell-type and tissue-specific gene expression (Fig. 4f). For example, subclass C17 was enriched for *Mef* family motifs, key regulators of muscle development. Subclass C16 showed strong enrichment for pioneering developmental factors like *ATF*, *NeuroG2* and *Sox10*, indicating its role in specifying neural lineages. Subclass C10 was defined by motifs for *NFkB* and *Hnf6b*, implicating its activity in immune responses and hepato-pancreatic cell types, respectively. Enhancers critical for stem cells and development were also clearly resolved, with subclass C6 containing motifs for *HNF6* and *Nanog*. Subclass C7 was characterized by *Hox* factor binding sites, which are essential for embryonic patterning. Similarly, sub-clusters C3/C4 were defined by motifs for *GLIS3* and *Nkx2*, transcription factors involved in pancreatic and thyroid development.

To quantitatively assess the tissue-specificity of the regulatory activities predicted by our model, we computed the Tau index for each sequence class. The Tau index is a standardized metric widely employed in genomics and transcriptomics to measure the degree of specialization in a given expression or activity profile. Tau index computed based on promoter and enhancer sequence classes revealed a highly significant divergence between the two main element types (SFig. 10b,c). Enhancers class exhibited higher median Tau index than promoters, confirming their dominant role as the drivers of cell-type-specific gene expression.

Comparing the tissue-specificity among Enhancer Sequence Class, we observed a strong enrichment of C11 and C30 consistent with the established biochemistry enrichment (SFig. 10c). However, C11 and C30 possessed hierarchical sub-cluster, therefore, we further dissected the enrichment of enhancer sub-clusters to reveal fine-grained distinct functional archetypes. Among the sub-clusters, C20 and C10 contributed most to the Enhancer activity of C11, representing the sequence basis of C11 core enhancer elements. Subclass C10 enriched H2BK12ac marks of H1 Derived Mesenchymal Stem Cells, was defined by motifs for *NFkB* and *Hnf6b*, implicating its activity in immune responses and hepato-pancreatic cell types. However, subclass C20 was architecturally and compositionally divergent, with H3K9me3, H3K27me3 histone marks enrichment, and was correspondingly enriched for *NFkB* and *Hnf6b*. For the sub-clusters of C30, subclass C17 contributed most to the Enhancer activity, with the enrichment of H3K36me3 histone marks, and *NFkB* and *Hnf6b* motifs.

To further validate the regulatory mechanisms underlying enhancer sequence class, we also compared their chromatin accessibility landscapes using single-cell ATAC-seq data (Fig. 4g). As expected, the chromatin accessibility of enhancers class exhibited significantly higher Tau index than promoters among cell types (Mann-Whitney U test, P-value < 2.72e-81). The results further supported that tissue-specific enhancers are selected in different cellular contexts by a diverse and rotating cast of distinct.

The fine-grained sub-clusters of the Enhancer sequence class reveals that the extensive heterogeneity of enhancers might be primarily derived by cell-type-specific activity, which could be defined by distinct signatures of underlying transcription factor (TF) binding motifs. Comparison between promoter and enhancer sequence classes further underscores enhancers as the primary drivers of tissue or cell-type specificity, which further precisely orchestrate spatiotemporal gene activity during development and cell lineage specification.

### Predict cell-type-specific gene expression from sequence

Accurately predicting gene expression from DNA sequence requires a model capable of decoding the complex grammar of the transcriptional regulation, including the function of promoters, enhancers, and their intricate interactions. Building upon the comprehensive context-aware sequence embeddings derived from HERMES, we constructed a specialized Transformer-based architecture to predict cell-type-specific gene expression levels (Fig. 5a). Leveraging the established vocabulary of sequence classes, we could further explain the deep-learning-based sequence-to-expression model, by systematically partitioning the genome based on sequence-intrinsic regulatory potential.

Specifically, we trained and validated a gene expression modeling framework using the ARCHS4^39^ database, which is a comprehensive compendium of 3,065 diverse biological states. By pairing gene regulatory sequence with expression profiles, we utilized HERMES to extract context-aware sequence embeddings, and then predict quantitative gene expression levels via a Transformer-based architecture. This gene-centred approach effectively aggregated cis-regulatory information into holistic expression levels spanning a wide array of cell type and tissue conditions, which also enabled precise alignment with downstream phenotypic data.

To evaluate the model’s predictive fidelity in reconstructing the global expression profiles of all held-out genes for a specific cellular condition, we first calculated the per-cell metric for each cell state. The model achieved compelling and highly accurate predictions, with a median per-cell Pearson correlation (PCC) of 0.83 (Fig. 5b) and a median per-cell Spearman correlation (SCC) of 0.82 (SFig. 11a) between the predicted and observed expression values across 3,065 cellular conditions. Per-cell accuracy exhibited a moderate positive correlation with overall transcriptional activity (Fig. 5b and SFig. 11a), suggesting that cell types with higher global transcriptional output were predicted with slightly greater fidelity. To compare our model’s performance in the context of existing methods, we also conducted a comprehensive benchmark against several state-of-the-art models for expression prediction. As shown in the benchmark analysis (SFig. 11c), our model demonstrated superior predictive performance. Specifically, our model achieved a higher median prediction accuracy and a tighter distribution of performance scores compared to other leading approaches. Notably, this performance closely approached the practical upper bound of predictability within the Arches dataset, which were 0.90 (for the mean expression) and 0.85 (for the median expression) based on the average correlation between consensus and individual tissue expression profiles.

Beyond evaluating the global gene expression prediction per cell, we rigorously evaluated the model’s ability to reconstruct the cell-type-specific expression patterns within each gene. The per-gene evaluation represents a more challenging task, which requires the model to resolve the differential gene expression patterns across diverse cellular contexts. HERMES demonstrated robust performance on per-gene metric, achieving a median PCC of 0.479 and a median SCC of 0.422 across 1,216 held-out genes (Fig. 5c). Notably, Per-gene PCC values remained remarkably correlated with expression abundance of genes (Fig. 5c), with performance attenuated for genes with extremely low expression. However, Per-gene SCC values remained largely independent of average gene expression levels, indicating that the model still preserved the correct relative rank order for low expressed genes (SFig. 11b).

Visual inspection of predicted and observed gene expression further illustrated the predictive fidelity of the model. As shown in representative tissues, predicted expression of held-out genes exhibited high correlations with experimentally measured gene expression across diverse cell conditions (Fig. 5d). Extending individual cellular state to the global scale, comparison of the full gene expression matrices confirmed that the model accurately reconstructed the complex landscape of transcriptional activity. Heatmaps of the predicted expression matrix faithfully recapitulated the structural patterns and intensity distributions of the target RNA-seq data (SFig. 12a,b), with the difference map confirming minimal systematic deviation between predicted and observed values (SFig. 12c). We further challenged the model to reproduce the higher-order co-expression relationships defining tissue-specific regulation. Comparison of gene-gene correlation matrices revealed that HERMES faithfully reconstructs the co-regulatory modules observed in the ground truth data with PCC of 0.272 (SFig. 13a). Quantitative assessment of cell-cell correlations also confirmed the tissue specificity of expression, yielding PCC of 0.743 between predicted and target cell-cell relationship pair (SFig. 13b).

Overall, our gene expression prediction model demonstrates a robust and multi-faceted ability to predict diverse cell-type-specific gene expression. The model not only achieves high accuracy in predicting global, cell-level expression profiles but also captures the more challenging nuances of tissue-specific gene expression, including the co-regulation patterns within gene networks. These results provide strong evidence that the model has successfully learned a generalizable mapping from sequence to cellular condition-specific expression states, reflecting a sophisticated understanding of transcriptional regulation in complex cellular environments.

### In silico enhancer perturbation reveal the logic of gene regulation

A primary goal in functional genomics is to move beyond expression prediction to an in silico framework that can mechanistically decipher gene regulation. This requires accurately identifying functional enhancers for a target gene and quantifying their regulatory effect in a specific cellular context. Having demonstrated our model’s predictive accuracy, we next sought to rigorously validate the model’s ability to prioritize enhancer-gene pairs.

We first benchmarked in silico contribution scores against experimental CRISPRi perturbation datasets^40,41^, by comparing the performance across different genomic distances and cell-type-specific scores. Our model accurately identified functional enhancer-gene pairs across diverse distances and cell types. Stratified by distance in a K562 cell-specific scenario, our scores showed robust performance for enhancers located in different distances from their target gene (Fig. 6a). Furthermore, we benchmarked the cell-type-specific contribution scores to evaluate the tissue-specific regulatory logic. We observed the same high performance across diverse cell lines, including HCT116, Jurkat, K562 and WTC11 (Fig. 6b). Critically, the cell-specific score was more accurate than other biochemical marks in the WTC11 cell line, providing direct evidence that the model has learned to utilize different sequence features in different cellular contexts. We also validated the model’s capability by dissecting the regulatory landscape of the PCBP1 gene (Fig. 6c). The perturbation prediction successfully prioritized the experimental CRISPRi effects, integrating the regulatory signals from different biochemical profiles, and further identified the cell-type-specific activity of enhancers.

Beyond validating our model on CRISPRi-tested enhancers, we next leveraged its predictive power to systematically quantify the effect size of all potential enhancers linked to the nearby genes. In total, the entire regulatory landscape of predicted enhancer effects covered 30,173 enhancers and 2,211 held-out genes across 3,065 cellular conditions. A subset of 7,689 enhancers were identified as high-specificity with a Tau index larger than 0.75, which revealed a highly dynamic regulatory landscape across tissues (SFig. 14a). Furthermore, correlation matrix of the predicted enhancer effects revealed distinct modules, with unsupervised clustering organizing tissues into hierarchically related biological lineages (SFig. 14b).

We then compared the enhancer perturbation effects between stable HKGs versus dynamic HVGs. The results showed a highly significant difference. Compared to HKGs, HVGs exhibited a significantly greater absolute mean enhancer effect (Mann-Whitney U test p<1e-8, Fig. 6d), and also a significantly higher variance of enhancer effect across all cell types (Mann-Whitney U test p<1e-8, Fig. 6e), indicating HVGs reliance on regulatory enhancers to drive expression across different cellular contexts.

In order to analyze the regulatory complexity across all the genes, we computed a Genomic Fragility Index to quantify the sensitivity of each gene to non-coding enhancer perturbations. The most fragile genes, which exhibit the high dependency on enhancers, were enriched for internal structural integrity and developmental precision, specifically extracellular matrix organization and vasculature development (SFig. 15a). In contrast, the most robust Genes, which utilize a promoter-dominant architecture to ensure high-output resilience, were enriched for external defense mechanisms, including antimicrobial humoral response and skin barrier formation (SFig. 15b).

Collectively, the in silico enhancer perturbation prediction provides compelling evidence that our model has learned the complex sequence-based grammar of enhancer-driven gene regulation. Our findings suggest that HVGs rely on regulatory enhancers to drive expression across diverse cellular contexts, and genes exhibiting high enhancer sensitivity evolved in the maintenance of multi-cellular and developmental complexity.

### Derived a formula of gene regulation from sequence-to-function model

While deep learning-based sequence-to-function models demonstrate remarkable predictive power, their inherent complexity often obscures the direct extraction of simple biological knowledge. Given that we have established HERMES to construct a comprehensive genome vocabulary of sequence classes and accurately predict tissue-specific gene expression, we then sought to distill its learned relationships into an interpretable mathematical formula.

To achieve this, we proposed a linear model that approximates the predictions of our complex HERMES model. This linear model was designed to represent gene expression as a function of core regulatory components, using the activities of promoters and enhancers as its primary variables, conditioned on a specific cellular context^29,42^. Crucially, we leveraged the gene expression prediction and enhancer perturbation prediction of the HERMES model as a causal reference to estimate the parameters of the linear model. The ultimate objective of this approach was to derive a formula of gene expression rule, which is an explicit and interpretable equation that quantifies the contribution of different cis-regulatory elements to gene expression in a context-specific manner.

To formalize the physical constraints governing the gene expression regulation dichotomy, we derived a unified kinematic equation that recapitulates the observed expression dynamics (Fig. 7a). We modeled the expression *T_g_*_,*c*_ of gene *g* in cellular context *c* as a function of its regulatory genomic sequence *S_g_*, decomposed into the linear coupling of intrinsic promoter strength (*P_g_*_,*c*_) and the cumulative enhancer activation (*E_i_*_,*g*,*c*_) from specific enhancer *i* link to gene *g* in tissue *c* within an insulator-delimited topological domain. Here, basal transcriptional frequency of *P_g_*_,*c*_ was denoted as *a*, and the enhancer contribution was modeled as coupling coefficient *b_i_*, which was determined by the intrinsic strength of the enhancer *E_i_*_,*g*,*c*_ and promoter *P_g_*_,*c*_.

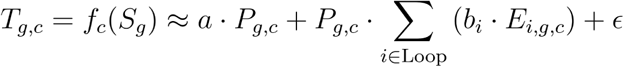

To quantify the causal contribution of individual regulatory elements, we exploited the HERMES model’s capacity for in silico perturbation. Unlike observational studies that rely on correlation, this framework allowed us to estimate causal regulatory effects by predicting expression ( 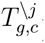) of a counterfactual sequence (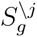) specifically lacking enhancer *j*. By approximating the deep neural network function *f_c_*^(*S*)^ with a linear model, we defined the expression in the absence of enhancer *j* as the baseline promoter activity coupled with the remaining enhancers.

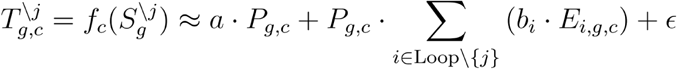

Consequently, the marginal contribution of the specific enhancer *j* was isolated by the difference between the full and perturbed states in our perturbation analysis. This derivation facilitated the linear model to capture the causal dependency of gene expression on specific enhancer-promoter interactions.

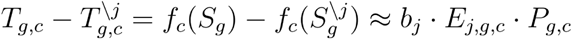

Finally, we estimated the parameters of the linear formula based on the sequence embeddings, tissue-specific gene expression prediction, and enhancer perturbation effect derived from HERMES. Specifically, by categorizing promoters and enhancers into distinct sequence classes, we constructed a pairwise interaction matrix **B** representing the intrinsic coupling efficiency *b*_(*P*,*E*)_between these regulatory classes (Fig. 7b).

Analysis of this interaction landscape revealed a highly structured cis-regulatory grammar rather than a random distribution of affinities. We observed striking functional specificity where certain promoter classes exhibit preferential compatibility with specific enhancer groups (Fig. 7b). For instance, Promoter related sequence classes of C7 and C19 demonstrated a robust activation module, forming strong positive couplings with a distinct set of enhancer classes (e.g., Enhancer related sequence classes of C11, C12, C13 and C21), suggesting a coordinated mechanism for high-output gene expression. Most notably, our model captured sophisticated context-dependent logic within the regulatory syntax. For example, Enhancer related sequence class C21, which acted as a strong activator for Promoter class C7 (b = +0.0172) yet switched to a potent repressor for Promoter class C5 (b = −0.0170). Promoter related sequence class C20 exhibited broad negative interaction across multiple enhancer types, yet functioned as a positive element for enhancer class C25 (b = −0.0087). This bipolar functionality underscored that the regulatory outcome of enhancer was not an inherent property of its sequence alone but was dictated by its compatibility with the cognate biochemical state of the promoter.

Synthesizing all the results, we propose a unified model of Enhancer-Promoter (E-P) compatibility that governs the distinct regulatory landscapes of the genome (Fig. 7c). HKGs predominantly utilize a strong-promoter architecture, enriched with ETS, YY1 and CCAAT motifs, forming strong positive couplings with enhancer classes to drive high-output expression across diverse cellular contexts. In contrast, HVGs relies on the context-dependent regulation of promoters and enhancers, characterized by underlying tissue or cell-type specific motifs, orchestrating spatiotemporal gene expression through a sophisticated regulatory syntax in a specific cellular context.

Ultimately, our approach bridges the gap between foundational predictive model and biological mechanistic understanding, by effectively transforming the latent representations of sequence-to-function model into a comprehensive functional vocabulary and a quantitative rulebook of transcription regulation.

## Discussion

In summary, by training HERMES on an unprecedented compendium of 137,127 functional genomics profiles, we established a foundational vocabulary of sequence classes across comprehensive genomic, functional and cellular contexts. Crucially, we leveraged the regulatory sequence to cell-type-specific gene expression, and distilled the deep learning model into a biologically interpretable transcriptional regulation formula. Our findings revealed the distinct promoters and motifs driving housekeeping genes, and quantitative rulebooks of Enhancer-Promoter (EP) compatibility underlying cell-type-specific highly variable genes.

A fundamental question in genomics is to understand the diverse function of invariant genomic sequence under dynamic cellular states. With trans-regulatory factors concentration fixed at a specific cellular context, the cis-regulatory elements encoded within the genome sequences respond specifically to trans-regulation. HERMES bridges the gap by hierarchical encoding complex genomic sequences into distinct regulatory mechanisms and expression syntax, using a sequence-to-function foundation model across the entire regulatory genome and functional profiles. The primary advantage of HERMES lies in its redefinition of the foundation model, which is not merely a predictive model, but also an interpretable atlas of cis-regulatory elements and the compatibility rules between enhancers and promoters.

HERMES unveiled the heterogeneity of regulatory sequences, resolving traditional broad categories into specific sequence classes. Our analysis resolved the monolithic promoter class into mechanistically distinct subtypes, including TATA-box dependent promoters, housekeeping-related promoters, and spatially regulated developmental promoters. Similarly, enhancer categories were also dissected into cell-type-specific subtypes defined by distinct underlying transcription factor binding motifs. High granular classification of regulatory elements derived from the sequence-to-function model shifts the paradigm from simple marks to a comprehensive functional vocabulary, revealing that the regulatory landscape of the genome sequence is more diverse and specialized than previously understood.

By integrating these regulatory sequence classes, we further derived a cis-regulatory formula that governs gene expression rules. Our analysis revealed that constitutively expressed genes (HKGs) exhibit a robust promoter-centric regulation, whereas highly variable genes (HVGs) are driven by weak promoters paired with tissue-specific enhancers. The regulatory architecture of genes suggests the compatibility rules for biochemically recruiting RNA Polymerase. The promoters of HKGs appear to encode high-affinity motifs that intrinsically stabilize the Pre-Initiation Complex, driving continuous Polymerase loading and transcription. Conversely, the weak promoters of HVGs rely on synergistic interactions with tissue-specific enhancers to construct a stable Transcriptional Complex under specific cellular conditions. The Interaction coefficient matrix of sequence classes also quantified the specific compatibility rules between promoters and enhancers, where the regulatory outcome is dictated by the precise matching of promoter strength with enhancer potential. Corroborating our findings, independent experimental studies have uncovered similar intrinsic compatibility rules by systematically permuting synthetic enhancers and promoters using STARR-seq technology^43,44^.

However, several limitations remain in our current framework. First, with our research objective set to describe the sequence class of native genome sequences, we trained the sequence-to-function model based on paired reference genome sequences and functional sequencing profiles, without any explicitly curated data from population mutations, induced perturbations and synthetic sequences. Although sequence-to-function models possess inherent zero-shot predictive capabilities, we did not evaluate the predictive performance of HERMES to quantify the impact of synthetic sequences and specific variants. Consequently, the sequence class vocabulary learned by HERMES might not be applicable for population mutation effect prediction, CRISPR-perturbated sequences or synthetic regulatory sequences with non-canonical motifs or novel combinations^45,46^. Second, while HERMES effectively compressed the vast landscape of functional profiles into a unified regulatory vocabulary, this reliance on existing data imposed inherent constraints. Therefore, the sequence vocabulary and expression syntax derived from the HERMES model were bounded by the biological diversity represented in our training atlas. For biological context, the model is not yet applicable to transient developmental intermediates or rare disease-associated configurations that lie significantly outside the distribution of the collected profiles. Furthermore, HERMES is also unable to predict unseen cellular conditions if those states are not explicitly captured in the underlying functional genomics data.

Looking forward, the HERMES framework is adaptable rather than being tethered to a specific neural architecture. The conceptual framework of HERMES is prioritizing the hierarchical encoding of functional states and the extraction of regulatory syntax. HERMES can seamlessly integrate more powerful sequence-to-function models, such as AlphaGenome, and emerging Genomic Language Models (GLMs), such as Evo^47^. Integrating more well-designed functional data or generative modeling approaches will also expand the framework’s predictive horizon to genuinely unmapped biological contexts and synthetic sequence vocabulary.

In summary, HERMES provides a unified and interpretable framework for analyzing the functional genome. By establishing the fundamental vocabulary of sequence classes and systematically decoding the combinatorial grammar of their interactions, HERMES serves as a foundational step towards the challenge of deciphering the complex language encoded in the genome sequence.

## Author contributions

J.L. implemented the packages, performed computational data analyses, and wrote the initial draft of the manuscript.

## Acknowledgments

The authors acknowledge that Gemini and Nano Banana (Google) was used for assistance with language editing and drew the illustration figures of the manuscript.

